# A broadly neutralizing antibody protects against SARS-CoV, pre-emergent bat CoVs, and SARS-CoV-2 variants in mice

**DOI:** 10.1101/2021.04.27.441655

**Authors:** David R. Martinez, Alexandra Schaefer, Sophie Gobeil, Dapeng Li, Gabriela De la Cruz, Robert Parks, Xiaozhi Lu, Maggie Barr, Kartik Manne, Katayoun Mansouri, Robert J. Edwards, Boyd Yount, Kara Anasti, Stephanie A. Montgomery, Shaunna Shen, Tongqing Zhou, Peter D. Kwong, Barney S. Graham, John R. Mascola, David. C. Montefiori, Munir Alam, Gregory D. Sempowski, Kevin Wiehe, Kevin O. Saunders, Priyamvada Acharya, Barton F. Haynes, Ralph S. Baric

**Affiliations:** Department of Epidemiology, University of North Carolina at Chapel Hill, Chapel Hill, NC, USA; Duke Human Vaccine Institute, Duke University School of Medicine, Durham, NC, USA; Lineberger Comprehensive Cancer Center, University of North Carolina School of Medicine, Chapel Hill, NC, USA; Department of Laboratory Medicine and Pathology, University of North Carolina School of Medicine, Chapel Hill, NC, USA; Vaccine Research Center, National Institute of Allergy and Infectious Diseases, NIH, Bethesda, MD, USA; Department of Surgery, Duke University School of Medicine, Durham, NC, USA

**Keywords:** SARS-CoV-2 D614G, B.1.1.7, B.1.429, B1.351, SARS-like virus, *Sarbecovirus*, countermeasures, DH1047 antibody, bNAb, panCoV

## Abstract

SARS-CoV in 2003, SARS-CoV-2 in 2019, and SARS-CoV-2 variants of concern (VOC) can cause deadly infections, underlining the importance of developing broadly effective countermeasures against Group 2B Sarbecoviruses, which could be key in the rapid prevention and mitigation of future zoonotic events. Here, we demonstrate the neutralization of SARS-CoV, bat CoVs WIV-1 and RsSHC014, and SARS-CoV-2 variants D614G, B.1.1.7, B.1.429, B1.351 by a receptor-binding domain (RBD)-specific antibody DH1047. Prophylactic and therapeutic treatment with DH1047 demonstrated protection against SARS-CoV, WIV-1, RsSHC014, and SARS-CoV-2 B1.351infection in mice. Binding and structural analysis showed high affinity binding of DH1047 to an epitope that is highly conserved among Sarbecoviruses. We conclude that DH1047 is a broadly neutralizing and protective antibody that can prevent infection and mitigate outbreaks caused by SARS-like strains and SARS-CoV-2 variants. Our results argue that the RBD conserved epitope bound by DH1047 is a rational target for pan Group 2B coronavirus vaccines.

## Introduction

The emergence of severe acute respiratory syndrome (SARS-CoV) in 2003 led to more than 8,000 infections and 800 deaths ^1,2^. In 2012, the Middle East Respiratory Syndrome (MERS-CoV) emerged in Saudi Arabia ^3^, which has so far infected ~2,600 people and caused 900 deaths. Less than a decade following the emergence of MERS-CoV, SARS-CoV-2 emerged in Wuhan, China ^4^. The spread of SARS-CoV-2, the virus that causes coronavirus disease of 2019 (COVID-19) was rapid, and by March 2020, the World Health Organization (WHO) had declared SARS-CoV-2 a global pandemic. By April 2021, more than 140 million people had been infected globally, resulting in >3 million deaths. Therefore, there is a need to develop safe and effective broad-spectrum countermeasures that can prevent the rapid spread and attenuate the severe disease outcomes associated with current and future SARS-like virus emergence events.

Human highly pathogenic CoV outbreaks are likely of bat origin ^5^, and there is great genetic diversity among bat SARS-like viruses ^6^. Zoonotic CoVs of bat origin, such as RsSHC014 and WIV-1, can utilize the human ACE2 receptor for cell entry and infect human airway cells ^7,8^, underlining their potential for emergence in naïve human populations. Moreover, existing SARS-CoV therapeutic monoclonal antibodies and SARS-CoV-2 mRNA vaccines do not protect against zoonotic SARS-like virus infection ^7–9^. Given the pandemic potential of SARS-like viruses, the development of broadly effective countermeasures, such as universal vaccination strategies ^9–11^, and coronavirus (CoV) cross-reactive monoclonal antibodies is a global health priority. Moreover, given the emergence of the SARS-CoV-2 variants that are partially or fully resistant to some neutralizing antibodies authorized for COVID-19 treatment ^12–14^, there is a need to discover mAb therapies that are broadly effective against the SARS-CoV-2 variants and zoonotic SARS-like viruses that will continue to emerge in the future.

The receptor binding domain (RBD) of SARS-CoV-2 is one of the targets for highly potent neutralizing antibodies. Despite the high degree of genetic diversity within the RBD in SARS-like viruses ^6^, antibodies can be engineered to recognize diverse SARS-like viruses. Rappazzo *et al.* recently reported that an engineered RBD-directed antibody, ADG-2, neutralized SARS-like viruses and protected against SARS-CoV and wild type SARS-CoV-2 ^15^. Therefore, the RBD of Sarbecoviruses contains conserved epitopes that are the target of broadly neutralizing antibodies. In agreement with the notion that the RBD contains a conserved epitope shared among SARS, SARS-like, SARS-CoV-2 and the variants, we have identified a panCoV protective antibody: DH1047. Here, we demonstrate, using both pseudoviruses and live virus assays, that DH1047 neutralizes SARS-CoV, SARS-like bat viruses RsSHC014 and WIV-1, and SARS-CoV-2 D614G, B.1.1.7, B.1.429, B.1.351 variants. Structural analysis shows that DH1047 targets a highly conserved RBD region among the Sarbecoviruses. Importantly, we also demonstrate that DH1047 provides prophylactic and therapeutic protection activity against pathogenic SARS-CoV, RsSHC014, WIV-1, wild type SARS-CoV-2, and against a pathogenic B.1.351 variant in mice. Thus, DH1047 is a pan-group 2B CoV protective antibody that can be used to prevent and treat SARS-CoV-2 infections including important with variants of concern and has the potential to prevent disease from a future outbreak of a pre-emergent, zoonotic SARS-like virus strains that jump into naïve animal and human populations.

## Results

### The identification of broadly cross-binding and neutralizing antibodies

We previously isolated 1737 monoclonal antibodies (mAbs) from a SARS-CoV convalescent patient 17 years following infection and a SARS-CoV-2 convalescent patient from 36 days post infection ^16^. From this large panel of mAbs previously described by Li *et al.* we focused on 50 cross-reactive antibodies which bound to SARS-CoV, SARS-CoV-2, and other human and animal CoV antigens ^16^. To examine if these cross-reactive mAbs neutralized divergent Sarbecoviruses, we measured neutralizing activity against a mouse-adapted SARS-CoV-2 2AA mouse-adapted (MA) virus, SARS-CoV, bat CoV WIV-1, and bat CoV RsSHC014 using live viruses, and found four broadly cross-reactive antibodies, DH1235, DH1073, DH1046, and DH1047 (Fig. 1). DH1235 neutralized SARS-CoV-2 2AA MA, SARS-CoV, and bat CoV WIV-1 with IC_50_ of 0.122, 0.0403, and 0.060 μg/ml, respectively (Fig. 1A and Table S1). DH1073 neutralized SARS-CoV-2 2AA MA, SARS-CoV, and bat CoV WIV-1 with IC_50_ of 0.808, 0.016, and 0.267 μg/ml, respectively (Fig. 1B and Table S1). DH1046 neutralized SARS-CoV-2 2AA MA, SARS-CoV, bat CoV WIV-1, and bat CoV RsSHC014 with IC_50_ of 2.85, 0.103, 0.425, and 1.27μg/ml, respectively (Fig. 1C and Table S1). Similar to DH1046, DH1047 more potently neutralized SARS-CoV-2 2AA MA, SARS-CoV, bat CoV WIV-1, and bat CoV RsSHC014 with IC_50_ of 0.397, 0.028, 0.191, and 0.200μg/ml, respectively (Fig. 1D and Table S1).

**Figure 1.**
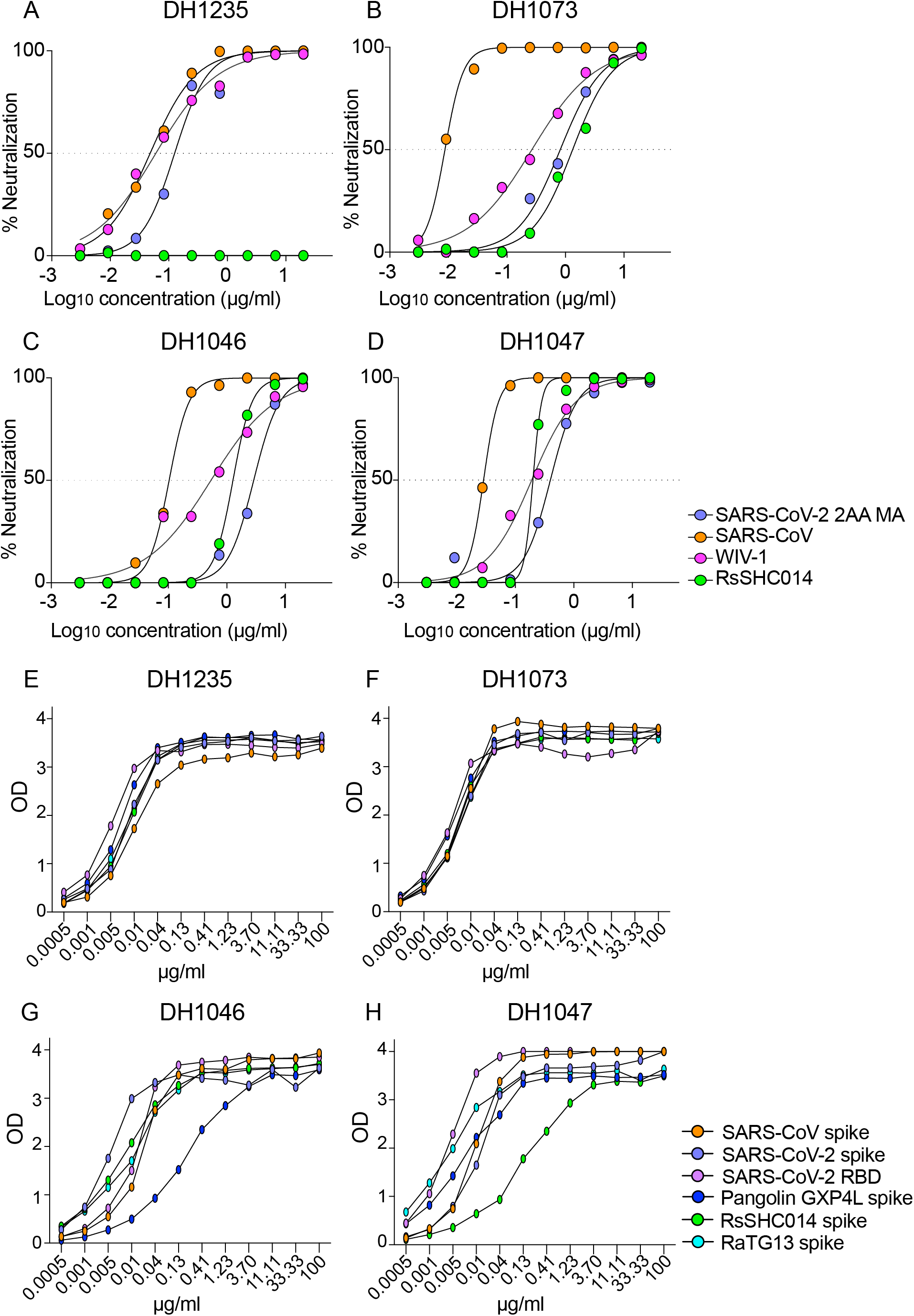
The identification of cross-reactive and broadly neutralizing antibodies. The neutralization activity of four broadly neutralizing antibodies against SARS-CoV-2 2AA mouse-adapted (MA), SARS-CoV, WIV-1, and RsSHC014. SARS-CoV-2 2AA MA is shown in purple, SARS-CoV is shown in orange, WIV-1 is shown in pink, and RsSHC014 is shown in green. The neutralization activity against Sarbecoviruses is shown for (A) DH1235, (B) DH1073, (C) DH1046, and (D) DH1047. The binding activity of cross-reactive antibodies against SARS-CoV spike, SARS-CoV-2 spike, SARS-CoV-2 RBD, Pangolin GXP4L spike, RaTG13 spike, and RsSHC014 spike of (E) DH1235, (F) DH1073, (G) DH1046, and (H) DH1047.

We also measured binding responses for DH1235, DH1073, DH1046, and DH1047 against zoonotic bat RaTG13-CoV, bat RsSHC014, and Pangolin GXP4L-CoV spikes. DH1235, DH1073, DH1046, and DH1047 mAbs showed strong binding to bat RaTG13-CoV, bat RsSHC014, and pangolin GXP4L-CoV spikes in addition to SARS-CoV and SARS-CoV-2 (Fig. 1E-1H). Finally, DH1235, DH1073, DH1046, and DH1047 bound to SARS-CoV-2 RBD and did not bind to the SARS-CoV-2 NTD, demonstrating specific binding to the RBD. While DH1235, DH1073, DH1046, and DH1047 were cross-reactive against epidemic, pandemic, and zoonotic Sarbecovirus spikes, they did not bind to MERS-CoV, HuCoV OC43, HuCoV NL63, and HuCoV 229E spike proteins (Fig. S1), suggesting these mAbs recognize a conserved epitope found only in Group 2B betacoronaviruses. By Negative Stain Electron Microscopy (NSEM), we observed binding of DH1047 to the RBD of bat RsSHC014 and SARS-CoV spike ectodomains, with overall similar orientations as was observed for DH1047 binding to the SARS-CoV-2 spike ectodomain (Figure S2) ^16^.

Finally, DH1235, DH1073, DH1046, and DH1047 exhibited medium to long heavy-chain-complementarity-determining-region 3 (HCDR3) lengths and variable somatic mutation rates in the heavy chain genes. DH1235, DH1073, and DH1046, had HCDR3 lengths of 21, 15, and 24, and somatic hypermutation (SMH) rates of 1.7, 9.0, and 4.7, respectively (Table S2). The most potent neutralizing antibody DH1047 had HCDR3 lengths and SMH rates of 24 and 8.05, respectively (Table S2).

### The protective activity of DH1235, DH1073, DH1046, and DH1047 against SARS-CoV

To define the protective efficacy of these four RBD-specific IgG bNAbs, we passively immunized aged mice with DH1235, DH1073, DH1046, DH1047 and a negative control influenza mAb, CH65 ^17^, at 10mg/kg 12 hours prior to infection and evaluated lung viral titer replication. Neither DH1235, DH1073, nor DH1046 protected against SARS-CoV mouse-adapted passage 15 (MA15) challenge in mice and all had lung viral replication comparable to that of control mice (Fig. 2A). In contrast, prophylactic administration of DH1047 fully protected mice from lung viral titer replication (Fig. 2A). Given the prophylactic potential of DH1047, we sought to also evaluate its therapeutic potential in a highly sensitive and stringent aged mouse model. We treated mice with control mAb and DH1047 at 10mg/kg both at 12 hours before and 12 hours post infection with SARS-CoV MA15 and monitored mice for signs of clinical disease, including weight loss, pulmonary function, which was measured by whole-body plethysmography (Buxco), through day 4 post infection (d4pi). In agreement with the SARS-CoV MA15 experiments, prophylactic treatment with DH1047 protected mice from weight loss through d4pi (Fig. 2B), and also protected mice from lung viral replication (Fig. 2C). We also evaluated if the prophylactic and therapeutic administration of DH1047 protected against lung pathology as measured by 1) lung discoloration, which is a visual metric of gross lung damage taken at the time of the necropsy, 2) microscopic evaluation as measured by an acute lung injury (ALI) scheme, and 3) a diffuse alveolar damage (DAD) scheme. ALI and DAD, which are characterized by histopathologic changes including alveolar septal thickening, protein exudate in the airspace, hyaline membrane formation, and neutrophils in the interstitium or alveolar sacs, were both blindly evaluated by a board-certified veterinary pathologist. The prophylactic administration of DH1047 resulted in complete protection from macroscopic lung discoloration (Fig. 2D) and microscopic lung pathology as measured by ALI (Fig. 2E and Fig. S3) and DAD (Fig. 2F and Fig. S3). Similarly, the therapeutic administration of DH1047 12 hours post infection resulted in reductions in lung viral titers (Fig. 2C and Fig. S3) as well as the macroscopic lung damage measured by the lung discoloration score (Fig. 2D and Fig. S3). In contrast to the prophylactic treatment condition, therapeutic administration of DH1047 did not significantly reduce microscopic lung pathology compared to control mice as measured by ALI (Fig. 2E and Fig. S3) and DAD (Fig. 2F and Fig. S3) in this highly susceptible model for SARS-CoV pathogenesis. Thus, DH1047 can prevent SARS-CoV disease when administered prophylactically and has early measurable therapeutic benefits in highly susceptible aged mouse models, much like other SARS-CoV-2 therapeutic neutralizing antibodies which have the most benefit in outpatient settings ^4,18,19^.

**Figure 2:**
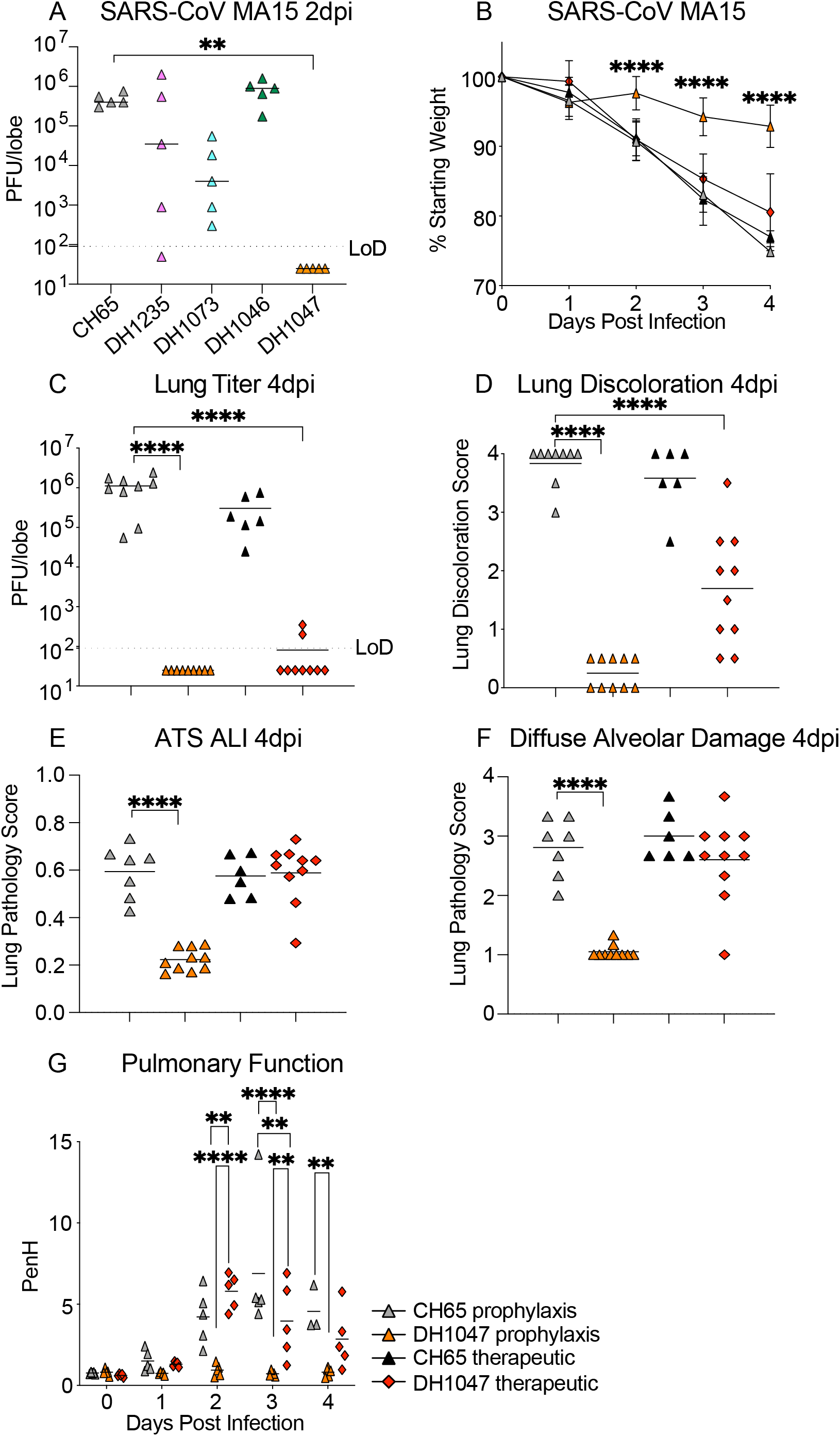
Prevention and therapy of DH1047 against SARS-CoV in aged mice. (A) SARS-CoV mouse-adapted 15 (MA15) lung viral replication in the prophylactically treated (−12 hours before infection) mice with a control influenza mAb CH65 and the four broadly neutralizing antibodies DH1235, DH1073, DH1046, DH1047. (B) % Starting weight of prophylactic (−12 hours before infection) and therapeutic (+12 hours after infection) treatment with DH1047 and control against SARS-CoV MA15 in mice. (C) Lung viral replication of SARS-CoV MA15 in mice treated prophylactically and therapeutically with DH1047 and control at 4 days post infection. (D) Macroscopic lung discoloration scores in mice treated with DH1047 and control prophylactically and therapeutically. (E) Lung pathology at day 4 post infection measured by acute lung injury (ALI) scores in mice treated with DH1047 and control prophylactically and therapeutically. (F) Lung pathology at day 4 post infection measured by diffuse alveolar damage (DAD) in mice treated prophylactically and therapeutically with DH1047 and control. (G) Pulmonary function as measured by whole body plethysmography (Buxco) in DH1047 and control mAb prophylactically and therapeutically treated mice. P values are from a 2-way ANOVA after Tukey’s multiple comparisons test for the weight loss, and P values are from a 1-way ANOVA following Dunnett’s multiple comparisons for the viral titer, and lung pathology readouts.

### Cryo-EM structure of the SARS-CoV/DH1047 complex

To visualize the binding epitope of DH1047 and to compare with the previously reported structure of the complex with the SARS-CoV-2 spike ectodomain^16^, we solved the cryo-EM structure of SARS-CoV bound to DH1047. 3D-classification of the cryo-EM dataset resulted in a 3.20 Å resolution reconstruction showing three DH1047 Fab bound to each of the 3 RBD of the ectodomain in the “up” position (1 Fab:1 RBD ratio) (Fig. 3, Figure S5 and Table S3). Similar to what we had observed for the DH1047 complex with the SARS-CoV-2 spike ectodomain ^16^, there was considerable heterogeneity in the RBD region; further classification of particles was performed to better resolve the antibody binding interface, resulting in an asymmetric reconstruction of a population refined to a resolution of 3.4 Å that was used for model fitting. The angle of approach and footprint of DH1047 on the SARS-CoV RBD closely resembled that in the SARS-CoV-2 complex with steric overlap predicted with ACE2 binding (Figure 3A-C). These results demonstrate that DH1047 binds to SARS-CoV and SARS-CoV-2 spike ectodomains by involving homologous interactions, consistent with our analysis of RBD sequence variability that showed a high degree of convergence of the DH1047 epitope ^10^, thereby defining an RBD conserved site of vulnerability among Sarbecoviruses. The DH1047 epitope on the SARS-CoV-2 RBD is distinct from other known antibodies of Classes 1, 2, 3 and 4 (Figure 3D). The DH1047 epitope overlays with that of antibody ADG-2, yet the two epitopes are distinct, and related by a rotation about the Fab longitudinal axis that pivots the ADG-2 antibody more towards the ACE2 binding region (Figures 3D and S6). Finally, we defined the binding affinity of DH1047 against epidemic and zoonotic spike proteins. We measured binding on and off rates against both SARS-CoV and RsSHC014-CoV spike proteins via surface plasmon resonance (SPR). DH1047 bound to the SARS-CoV and RsSCH014-CoV spikes with high affinity, association rates (> 8.60 X10^4^ M^−1^s^−1^) and dissociation rates (< 1.0X10^−5^ s^−1^) (Fig. S4), demonstrating that DH1047 binds tightly to both the epidemic SARS-CoV and pre-emergent bat CoVs that are poised for human emergence.

**Figure 3.**
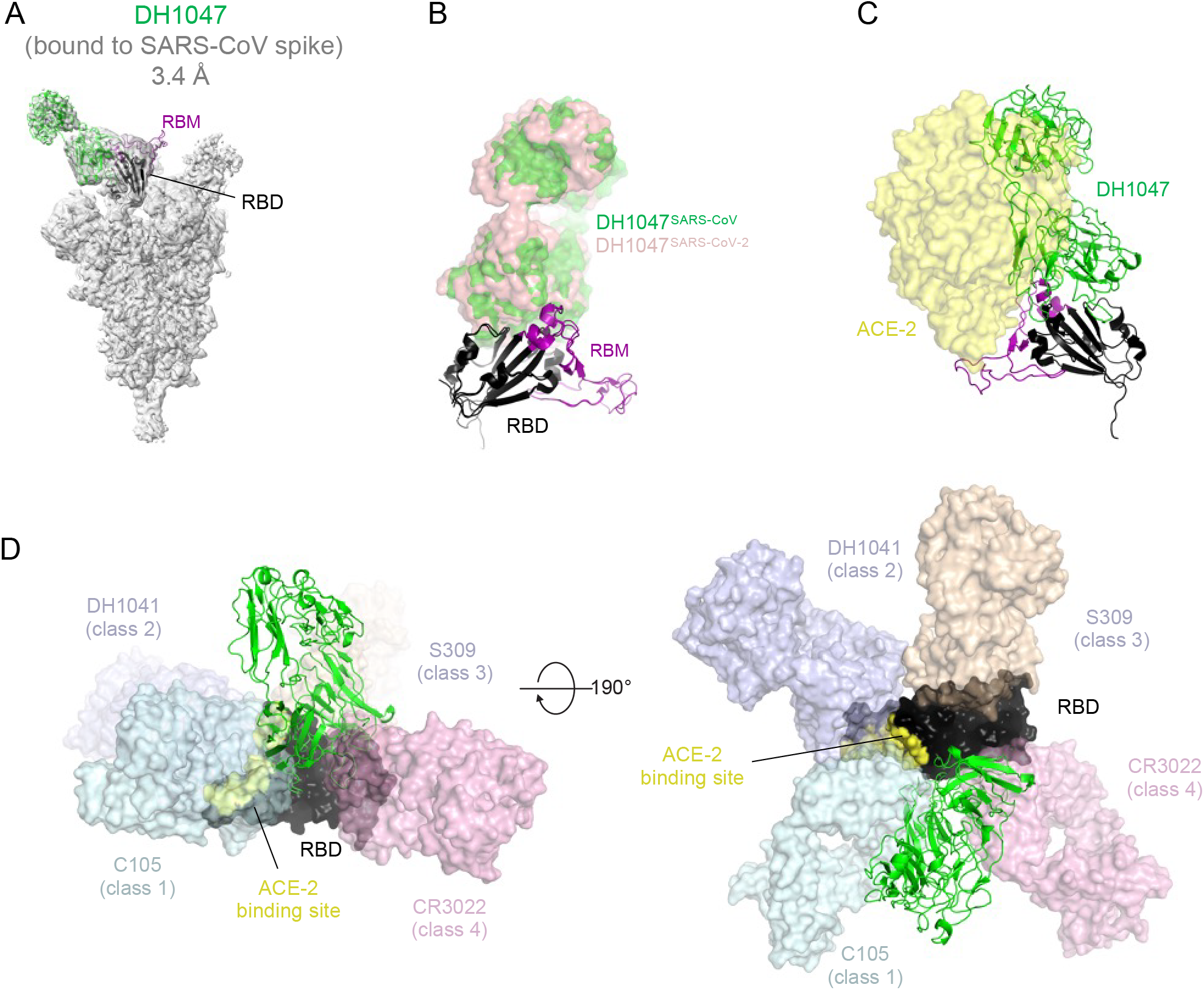
Cryo-EM structure of DH1047 bound to SARS-CoV spike. (A) Cryo-EM reconstruction of DH1047 Fab bound to SARS-CoV spike shown in grey, with the underlying fitted model shown in cartoon representation. DH1047 is colored green, the RBD it is bound to is colored black with the Receptor Binding Motif within the RBD colored purple. (B) Overlay of DH1047 bound to SARS-CoV-1 and SARS-CoV-2 (PDB ID: 7LDI) S proteins. Overlay was performed with the respective RBDs. DH1047 bound to SARS-CoV and SARS-CoV-2 spike is shown in green and and salmon, respectively. (C) ACE2 (yellow surface representation, PDB 6VW1) binding to RBD is sterically hindered by DH1047. The views in panels B and C are related by a ~180º rotation about the vertical axis. (D) DH1047 binding relative to binding of other known antibody classes that bind the RBD. RBD is shown in black with the ACE2 footprint on the RBD colored yellow. DH1047 is shown in cartoon representation and colored green. The other antibodies and shown as transparent surfaces: C105 (pale cyan, Class 1, PDB ID: 6XCN and 6XCA), DH1041 (light blue, Class 2, PDB ID: 7LAA), S309 (wheat, Class 3, PDB ID:6WS6 and 6WPT) and CR3022 (pink, Class 4, PDB ID: 6YLA)

### The prophylactic and therapeutic activity of DH1047 against bat pre-emergent CoVs and *in vitro* neutralization activity against the SARS-CoV-2 variants

As DH1047 neutralized both the pre-emergent bat CoVs WIV-1 and RsSHC014 (Fig. 1), we sought to define if DH1047 had prophylactic and therapeutic efficacy in mice. We evaluated the protective efficacy against lung viral replication against these pre-emergent bat CoVs. We administered DH1047 prophylactically 12 hours before infection and therapeutically 12 hours post infection at 10mg/kg in mice infected with bat CoVs. Importantly, the prophylactic administration of DH1047 completely protected mice from WIV-1 lung viral replication and reduced lung viral titers in therapeutically treated mice compared to control mice (Fig. 4A). Similarly, the prophylactic administration of DH1047 completely protected mice from RsSHC014 lung viral replication and significantly reduced viral replication to near undetectable levels in therapeutically treated mice (Fig. 4B). While we previously demonstrated the prophylactic and therapeutic efficacy of DH1047 against the wild type SARS-CoV-2 in cynomolgus macaques ^16^, which exhibit mild SARS-CoV-2 disease ^20^, it was not known if the mutations present in the newly emerging SARS-CoV-2 variants would ablate the neutralizing activity of DH1047. We therefore evaluated if DH1047 could neutralize the prevalent variants of concern (VOCs): SARS-CoV-2 D614G, SARS-CoV-2 UK B.1.1.7., SARS-CoV-2 California B1.429, and SARS-CoV South Africa B1.351 using both pseudovirus and live virus neutralization assays. DH1047 neutralized all tested variants of concern with substantial potency (Fig. 4C and Fig. 4D). Pseudovirus neutralization assays revealed strong neutralization of DH1047 against the SARS-CoV-2 VOCs (Fig. 4D). Importantly, live virus neutralization also demonstrated the broadly neutralizing activity of DH1047 with IC_50_ values against D614G, B.1.1.7, and B1.351 were 0.059, 0.081, and 0.111μg/ml, respectively.

**Figure 4:**
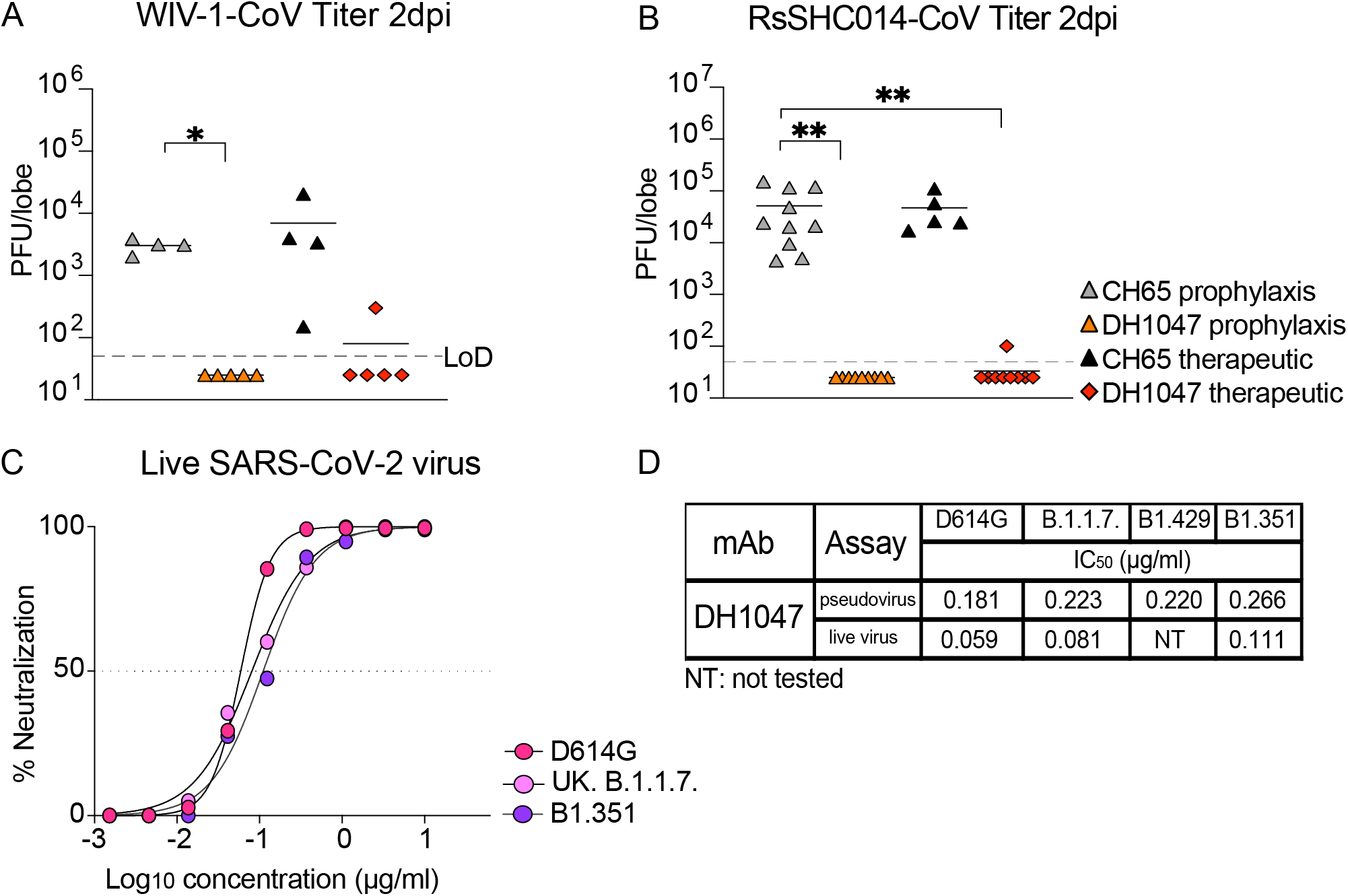
Prophylactic and therapeutic activity of DH1047 against SARS-like bat CoVs and the *in vitro* neutralization against the SARS-CoV-2 variants. Lung viral replication of WIV-1 in mice treated prophylactically and therapeutically with DH1047 and control at 2 days post infection. (B) Lung viral replication of RsSHC014 in mice treated prophylactically and therapeutically with DH1047 and control at 2 days post infection. (C) Live virus neutralization of SARS-CoV-2 D614G, UK B.1.1.7., and South African B.1.351 variants. (D) The comparison of the DH1047 neutralization activity against the SARS-CoV-2 variants in pseudovirus and live virus neutralization assays. P values are from a 2-way ANOVA after Tukey’s multiple comparisons test for the weight loss, and P values are from a 1-way ANOVA following Dunnett’s multiple comparisons for the viral titer, and lung pathology readouts.

### The prophylactic and therapeutic activity of DH1047 against SARS-CoV-2 B.1.351 in mice

Given that the B.1.351 South African variant is more resistant to both vaccine-elicited neutralizing antibodies ^14,21^, and completely ablates the neutralizing activity of the Eli Lily therapeutic monoclonal antibody LY-CoV555 ^12^, we also sought to evaluate if DH1047 had both prophylactic and therapeutic efficacy against SARS-CoV-2 B.1.351, which incorporates the B.1.351 spike in the SARS-CoV MA10 genome backbone ^22^. We again utilized a highly susceptible and vulnerable aged mouse model in the SARS-CoV-2 B.1.351 protection experiments. Consistent with the SARS-CoV, WIV-1, and RsSHC014 *in vivo* data, the prophylactic administration of DH1047 mediated protection against severe weight loss following SARS-CoV-2 B.1.351 challenge in aged mice (Fig. 5A). In contrast, we did not observe differences in weight loss from the therapeutic administration of DH1047 (Fig. 5A). Mice prophylactically treated with DH1047 had undetectable levels of SARS-CoV-2 B.1.351 lung viral replication (Fig. 5B) and were also completely protected from macroscopic lung pathology compared to controls (Fig. 5C). While we observed no significant protection from weight loss in DH1047 therapeutically treated mice, we did observe a significant reduction in lung viral titers compared to control (Fig. 5B). We also evaluated the microscopic lung pathology as measured by ALI (Fig. 5D) and DAD scoring schemes (Fig. 5E) in this highly susceptible aged model for SARS-CoV-2 B.1.351 pathogenesis. Importantly, the prophylactic administration of DH1047 significantly protected mice from lung histopathology as measured by ALI and DAD compared to control mice. Additionally, we observed a reduction in ALI by the therapeutic administration of DH1047 as measure by macroscopic lung pathology (Fig. 5C) and lung histopathology by ALI (Fig. 5D). Therefore, DH1047 can prevent and treat SARS-CoV-2 infections with the B.1.351 variant of concern *in vivo*, especially if given early in infection.

**Figure 5:**
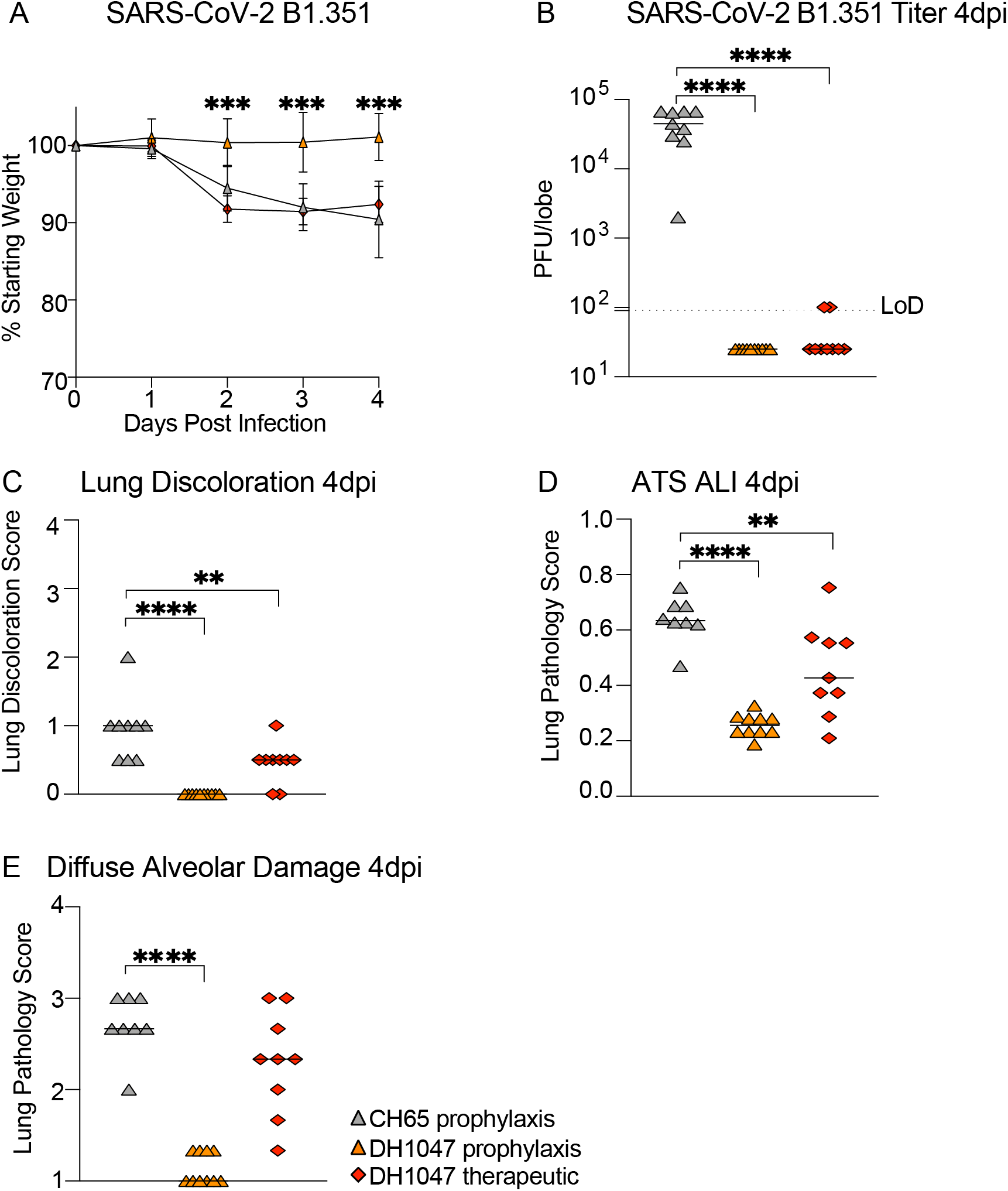
Prevention and therapy of DH1047 against SARS-CoV-2 B.1.351 in mice. (A) % Starting weight of prophylactic (−12 hours before infection) and therapeutic (+12 hours after infection) treatment with DH1047 and control against SARS-CoV-2 B.1.351 in mice. (B) Lung viral replication of SARS-CoV-2 B.1.351 in mice treated prophylactically and therapeutically with DH1047 and control at 4 days post infection. (C) Macroscopic lung discoloration scores in mice treated with DH1047 and control prophylactically and therapeutically. (E) Lung pathology at day 4 post infection measured by acute lung injury (ALI) scores in mice treated with DH1047 and control prophylactically and therapeutically. (F) Lung pathology at day 4 post infection measured by diffuse alveolar damage (DAD) in mice treated prophylactically and therapeutically with DH1047 and control. P values are from a 2-way ANOVA after Tukey’s multiple comparisons test for the weight loss, and P values are from a 1-way ANOVA following Dunnett’s multiple comparisons for the viral titer, and lung pathology readouts.

## Discussion

The emergence of SARS-CoV and SARS-CoV-2 in the last two decades underscores a critical need to develop broadly effective countermeasures against Sarbecoviruses. Moreover, with the recent emergence of more highly transmissible ^23^, virulent ^24^, and neutralization resistant UK B.1.1.7 variant, that can partially evade existing countermeasures ^12,14^, there is a need to develop next-generation mAb therapeutics that can broadly neutralize these variants, as well as future variants of concern. For example, the SARS-CoV-2 South African B.1.351 variant completely ablates the neutralization activity of the mAb LY-CoV555 ^12,13^. As a result, the emergency use authorization (EUA) of LY-CoV555 was recently rescinded by the U.S. Food and Drug Administration (FDA). In addition, the presence of the E484K mutation in many variants of concern, severely dampens the neutralization activity by more than 6-fold of the AstraZeneca COV2-2196 mAb, Brii BioSciences mAb Brii-198, and the Regeneron mAb REGN 10933 ^13,14,19^. In addition to evading currently monoclonal antibody therapeutics, some of the variants including B.1.351 can diminish the efficacy of clinically approved vaccines, including the Johnson & Johnson single-dose vaccine and the AstraZeneca ChAdOx1 ^25,26^. Furthermore, some monoclonal antibodies isolated from vaccine recipients of the Moderna and Pfizer vaccines also demonstrated reduced efficacy against mutations present in the variants ^27^. Therefore, current vaccine and mAb therapies must be monitored in real time to define the performance of existing therapies against newly emerging and spreading variants. In the setting of reduced vaccine efficacy, the deployment of effective mAb therapies against the variants, such as DH1047, could be a strategy to help control the COVID-19 pandemic.

The development of universal vaccination strategies against Sarbecoviruses will be improved by the identification and characterization of broadly protective and conserved epitopes across SARS-like virus strains. Recent studies described broadly reactive antibodies that target the subunit 2 (S2) portion of the spike protein ^28–31^. While the broad recognition of these S2-specific antibodies is encouraging, these antibodies weakly neutralized diverse CoVs. Given the limited characterization of these mAbs, it is unclear if these S2-specific mAbs are broadly protective *in vivo* against diverse epidemic and zoonotic pre-emergent CoVs. In contrast, RBD-specific antibody, S2X259, neutralized SARS-CoV-2 variants and zoonotic SARS-like viruses, as measured by pseudovirus neutralization^32^. Similarly, a recent subset of RBD-specific cross-reactive mAbs also showed *in vitro* activity ^33^, although their *in vivo* breadth and protective efficacy remains unconfirmed. It is interesting that DH1235, DH1073, and DH1046 neutralized SARS-CoV but did not protect against SARS-CoV challenge *in vivo.* Perhaps DH1235, DH1073, and DH1046 require 1) non-neutralizing functions for protecting against infection *in vivo*, or 2) have a distinct mode or angle of binding to SARS-CoV compared to DH1047 required for the observed protection. This underlines the importance of performing *in vivo* protection studies in addition to *in vitro* neutralization assays to truly define the protective efficacy of panCoV-specific mAbs.

In contrast to ADG-2 which uses VH3-21 for its heavy chain and has a 17 amino acid long HCDR3, DH1047 uses VH1-46 and has a 24 amino acid long HCDR3 (Table S2) ^15^. Moreover, ADG-2 and DH1047 have overlapping, yet distinct binding footprints, targeting a conserved region on the RBD (Fig. S6). In addition, the DH1047 epitope is distinct to those from cross-reactive antibodies S309 and CR3022 (Fig. 3D) and targets an epitope near those from class 4 antibodies. DH1047 had broad protective *in vivo* efficacy against pre-emergent SARS-like viruses, epidemic SARS-CoV, and the SARS-CoV-2 B.1.351 variant, underscoring that DH1047 recognizes a pan Sarbecovirus neutralizing epitope. Consistent with this notion, we have described a SARS-CoV-2 RBD-ferritin nanoparticle vaccine that elicited neutralizing antibodies against pre-emergent SARS-like viruses and protected against SARS-CoV-2 challenge in monkeys ^10^. The serum antibody responses in these SARS-CoV-2 RBD-ferritin nanoparticle-vaccinated monkeys could block DH1047 binding responses against SARS-CoV-2 spike proteins, suggesting that SARS-CoV-2 RBD vaccines elicit DH1047-like antibody responses and could potentially protect against the future emergence of SARS- or SARS2-like viruses.

Moving forward, it will be critical to closely monitor SARS- and SARS2-like viruses of zoonotic origin and actively monitor if broad-spectrum antibodies like ADG-2, DH1047, and S2X259 retain their inhibitory activity against pre-emergent viruses. We envision a system in which broad-spectrum antibodies like DH1047 could be tested for safety in small Phase I clinical trials so that in the event that a future SARS-like virus emerges, DH1047 could immediately be tested in larger efficacy trials at the site of an outbreak to potentially prevent the rapid spread of an emergent CoV. Moreover, given that DH1047 exhibited strong *in vivo* protection against the SARS-CoV-2 B.1.351 VOC, this mAb could be deployed as a mAb therapeutic to help control the current COVID-19 pandemic. Like other therapeutic antibodies evaluated against COVID19 infections, our data argues that early administration will prove critical for protecting against severe disease outcomes ^19^. We conclude that DH1047 is a broadly protective mAb that has efficacy against pre-emergent, zoonotic SARS-like viruses from different clades, neutralizes highly transmissible SARS-CoV-2 variants, and protects against SARS-CoV-2 B.1.351.

## Methods

### Antibody isolation

Antibodies were isolated from antigen-specific single B cells as previously described from an individual who had recovered from SARS-CoV-1 infection 17 years prior to leukapheresis, and from a SARS-CoV-2 convalescent individual from 36 days post infection ^16^.

### Measurement of CoV spike binding by ELISA

Indirect binding ELISAs were conducted in 384 well ELISA plates (Costar #3700) coated with 2μg/ml antigen in 0.1M sodium bicarbonate overnight at 4°C, washed and blocked with assay diluent (1XPBS containing 4% (w/v) whey protein/ 15% Normal Goat Serum/ 0.5% Tween-20/ 0.05% Sodium Azide). mAbs were incubated for 60 minutes in three-fold serial dilutions beginning at 100μg/ml followed by washing with PBS/0.1% Tween-20. HRP conjugated goat anti-mouse IgG secondary antibody (SouthernBiotech 1030-05) was diluted to 1:10,000 in assay diluent without azide, incubated at for 1 hour at room temperature, washed and detected with 20μl SureBlue Reserve (KPL 53-00-03) for 15 minutes. Reactions were stopped via the addition of 20μl HCL stop solution. Plates were read at 450nm. Area under the curve (AUC) measurements were determined from binding of serial dilutions.

### Measurement of neutralizing antibodies against live viruses

Full-length SARS-CoV-2 Seattle, SARS-CoV-2 D614G, SARS-CoV-2 B.1.351, SARS-CoV-2 B.1.1.7, SARS-CoV, WIV-1, and RsSHC014 viruses were designed to express nanoluciferase (nLuc) and were recovered via reverse genetics as described previously ^16^. Virus titers were measured in Vero E6 USAMRIID cells, as defined by plaque forming units (PFU) per ml, in a 6-well plate format in quadruplicate biological replicates for accuracy. For the 96-well neutralization assay, Vero E6 USAMRID cells were plated at 20,000 cells per well the day prior in clear bottom black walled plates. Cells were inspected to ensure confluency on the day of assay. mAbs were serially diluted 3-fold up to nine dilution spots at specified concentrations. Serially diluted mAbs were mixed in equal volume with diluted virus. Antibody-virus and virus only mixtures were then incubated at 37°C with 5% CO_2_ for one hour. Following incubation, serially diluted mAbs and virus only controls were added in duplicate to the cells at 75 PFU at 37°C with 5% CO_2_. After 24 hours, cells were lysed, and luciferase activity was measured via Nano-Glo Luciferase Assay System (Promega) according to the manufacturer specifications. Luminescence was measured by a Spectramax M3 plate reader (Molecular Devices, San Jose, CA). Virus neutralization titers were defined as the sample dilution at which a 50% reduction in RLU was observed relative to the average of the virus control wells.

### Surface plasmon resonance

Kinetic measurements of the DH1047 Fab binding to SARS-CoV and RsSHC014 spike proteins were obtained using a Biacore S200 instrument (Cytiva, formerly GE Healthcare) in HBS-EP+ 1X running buffer. The spike proteins were first captured onto a Series S Streptavidin chip to a level of 300-400 for the SARS-CoV spike proteins and 850-1000RU for the RsSHC014 spike protein. The DH1047 Fab was diluted from 2.5 to 200nM and injected over the captured CoV spike proteins using the single cycle kinetics injection type at a flow rate of 50μL/min. There were five 120s injections of the Fab at increasing concentrations followed by a dissociation of 600s after the final injection. After dissociation, the spike proteins were regenerated from the streptavidin surface using a 30s pulse of Glycine pH1.5. Results were analyzed using the Biacore S200 Evaluation software (Cytiva). A blank streptavidin surface along with blank buffer binding were used for double reference subtraction to account for non-specific protein binding and signal drift. Subsequent curve fitting analyses were performed using a 1:1 Langmuir model with a local Rmax for the DH1047 Fab. The reported binding curves are representative of 2 data sets.

### Protein expression and purification for EM studies

The SARS-CoV spike ectodomain construct comprised the residues 1 to 1190 (UniProt P59594-1) with proline substitutions at 968-969, a C-terminal T4 fibritin trimerization motif, a C-terminal HRV3C protease cleavage site, a TwinStrepTag and an 8XHisTag. The construct was cloned into the mammalian expression vector pαH^34^. The RsSHC014 spike ectodomain construct was prepared similarly, except it also contained the 2P mutations that placed two consecutive proline at the HR1-CH junction at residue positions 986 and 987. FreeStyle 293F cells were used for the spike ectodomain production. Cells were maintained in FreeStyle 293 Expression Medium (Gibco) at 37°C and 9% CO_2_, with agitation at 120 rpm in a 75% humidified atmosphere. Transfections were performed as previously described ^35–38^ using Turbo293 (SpeedBiosystems). 16 to 18 hours post transfection, HyClone CDM4HEK293 media (Cytiva, MA) was added. On the 6^th^ day post transfection, spike ectodomain was harvested from the concentrated supernatant. The purification was performed using StrepTactin resin (IBA LifeSciences) and size exclusion chromatography (SEC) on a Superose 6 10/300 GL Increase column (Cytiva, MA) in 2mM Tris, pH 8.0, 200 mM NaCl, 0.02% NaN_3_. All steps were performed at room temperature and the purified spike proteins were concentrated to 1-5 mg/ml, flash frozen in liquid nitrogen and stored at −80 °C until further use.

DH1047 IgG was produced in Expi293F cells maintained in Expi293 Expression Medium (Gibco) at 37°C, 120 rpm, 8% CO_2_ and 75% humidity. Plasmids were transfected using the ExpiFectamine 293 Transfection Kit and protocol (Gibco) ^35–37^ and purified by Protein A affinity. The IgG was digested to the Fab state using LysC.

### Negative Stain Electron Microscopy (NSEM)

NSEM was performed as described previously ^16^. Briefly, Fab-spike complexes were prepared by mixing Fab and spike to give a 9:1molar ratio of Fab to spike. Following a 1-hr incubation for 1 hour at 37 °C, the complex was cross-linked by diluting to a final spike concentration of 0.1 mg/ml into room-temperature buffer containing 150 mM NaCl, 20 mM HEPES pH 7.4, 5% glycerol, and 7.5 mM glutaraldehyde and incubating for 5 minutes. Excess glutaraldehyde was quenched by adding sufficient 1 M Tris pH 7.4 stock to give a final concentration of 75 mM Tris and incubated for 5 minutes. Carbon-coated grids (EMS, CF300-cu-UL) were glow-discharged for 20s at 15 mA, after which a 5-μl drop of quenched sample was incubated on the grid for 10-15 s, blotted, and then stained with 2% uranyl formate. After air drying grids were imaged with a Philips EM420 electron microscope operated at 120 kV, at 82,000x magnification and images captured with a 2k x 2k CCD camera at a pixel size of 4.02 Å.

The RELION 3.0 program was used for all negative stain image processing. Images were imported, CTF-corrected with CTFFIND, and particles were picked using a spike template from previous 2D class averages of spike alone. Extracted particle stacks were subjected to 2-3 rounds of 2D class averaging and selection to discard junk particles and background picks. Cleaned particle stacks were then subjected to 3D classification using a starting model created from a bare spike model, PDB 6vsb, low-pass filtered to 30 Å. Classes that showed clearly defined Fabs were selected for final refinements followed by automatic filtering and B-factor sharpening with the default Relion post-processing parameters.

### Cryo-EM

Purified SARS-CoV-1 spike ectodomain was incubated for approximatively 2 hours with a 6-fold molar equivalent of the DH1047 Fab in a final volume of 10μL. The sample concentration was adjusted to ~1.5 mg/mL of spike in 2 mM Tris pH 8.0, 200 mM NaCl, and 0.02% NaN_3_. Before freezing, 0.1μL of glycerol was added to the 10μL of sample. A 2.4-μL drop of protein was deposited on a Quantifoil-1.2/1.3 grid (Electron Microscopy Sciences, PA) that had been glow discharged for 10 seconds using a PELCO easiGlow™ Glow Discharge Cleaning System. After a 30-second incubation in >95% humidity, excess protein was blotted away for 2.5 seconds before being plunge frozen into liquid ethane using a Leica EM GP2 plunge freezer (Leica Microsystems). Frozen grids were imaged using a Titan Krios (Thermo Fisher) equipped with a K3 detector (Gatan). Data processing was performed using cryoSPARC ^39^. Model building and refinement was done using Phenix ^40,41^, Coot ^42^, Pymol ^43^, Chimera ^44^, ChimeraX ^45^ and Isolde ^46^.

### Animals and challenge viruses

Eleven-month-old female BALB/c mice were purchased from Envigo (#047) and were used for the SARS-CoV, SARS-CoV-2 B1.351, and RsSHC014-CoV protection experiments. 8-10-week-old hACE2-transgenic mice were bred at UNC Chapel Hill and were used for WIV-1-CoV protection experiments. The study was carried out in accordance with the recommendations for care and use of animals by the Office of Laboratory Animal Welfare (OLAW), National Institutes of Health and the Institutional Animal Care and Use Committee (IACUC) of University of North Carolina (UNC permit no. A-3410-01). Animals were housed in groups of five and fed standard chow diets. Virus inoculations were performed under anesthesia and all efforts were made to minimize animal suffering. All mice were anesthetized and infected intranasally with 1 × 10^4^ PFU/ml of SARS-CoV MA15, 1 × 10^4^ PFU/ml of SARS-CoV-2 B1.351-MA10, 1 × 10^4^ PFU/ml RsSHC014, 1 × 10^4^ PFU/ml WIV-1, which have been described previously ^7,22,47^. Mice were weighted daily and monitored for signs of clinical disease, and selected groups were subjected to daily whole-body plethysmography. For all mouse studies, groups of n=10 mice were included per arm of the study except for the hACE2-transgenic mice, which included n=5 mice per group due to a limited availability of these mice. Viral titers, weight loss, and histology were measured from individual mice per group.

### Lung pathology scoring

Acute lung injury was quantified via two separate lung pathology scoring scales: Matute-Bello and Diffuse Alveolar Damage (DAD) scoring systems. Analyses and scoring were performed by a board vertified veterinary pathologist who was blinded to the treatment groups as described previously ^48^. Lung pathology slides were read and scored at 600X total magnification.

The lung injury scoring system used is from the American Thoracic Society (Matute-Bello) in order to help quantitate histological features of ALI observed in mouse models to relate this injury to human settings. In a blinded manner, three random fields of lung tissue were chosen and scored for the following: (A) neutrophils in the alveolar space (none = 0, 1–5 cells = 1, > 5 cells = 2), (B) neutrophils in the interstitial septa (none = 0, 1–5 cells = 1, > 5 cells = 2), (C) hyaline membranes (none = 0, one membrane = 1, > 1 membrane = 2), (D) Proteinaceous debris in air spaces (none = 0, one instance = 1, > 1 instance = 2), (E) alveolar septal thickening (< 2x mock thickness = 0, 2–4x mock thickness = 1, > 4x mock thickness = 2). To obtain a lung injury score per field, A–E scores were put into the following formula score = [(20x A) + (14 x B) + (7 x C) + (7 x D) + (2 x E)]/100. This formula contains multipliers that assign varying levels of importance for each phenotype of the disease state. The scores for the three fields per mouse were averaged to obtain a final score ranging from 0 to and including 1. The second histology scoring scale to quantify acute lung injury was adopted from a lung pathology scoring system from lung RSV infection in mice ^49^. This lung histology scoring scale measures diffuse alveolar damage (DAD). Similar to the implementation of the ATS histology scoring scale, three random fields of lung tissue were scored for the following in a blinded manner: 1= absence of cellular sloughing and necrosis, 2=Uncommon solitary cell sloughing and necrosis (1–2 foci/field), 3=multifocal (3+foci) cellular sloughing and necrosis with uncommon septal wall hyalinization, or 4=multifocal (>75% of field) cellular sloughing and necrosis with common and/or prominent hyaline membranes. The scores for the three fields per mouse were averaged to get a final DAD score per mouse. The microscope images were generated using an Olympus Bx43 light microscope and CellSense Entry v3.1 software.

### Biocontainment and biosafety

Studies were approved by the UNC Institutional Biosafety Committee approved by animal and experimental protocols in the Baric laboratory. All work described here was performed with approved standard operating procedures for SARS-CoV-2 in a biosafety level 3 (BSL-3) facility conforming to requirements recommended in the Microbiological and Biomedical Laboratories, by the U.S. Department of Health and Human Service, the U.S. Public Health Service, and the U.S. Center for Disease Control and Prevention (CDC), and the National Institutes of Health (NIH).

### Statistics

All statistical analyses were performed using GraphPad Prism 9.

## Data availability

Structural data of DH1047 will be made available after publication.

## Code availability

No code was generated in this study.

## Author contributions

Conceived the study: D.R.M., A.S., S.G., D.L., P.A., B.F.H., R.S.B. designed experiments: D.R.M., A.S., S.G., D.L., performed laboratory experiments: D.R.M, A.S., D.L., S.G., Provided critical reagents: T.Z., P.D.K., B.S.G., and K.O.S. Analyzed data and provided critical insight: D.R.M, A.S., S.G., D.L., G.D.LC., R.P., M.B., K.M., B.Y., K.A., S.M., T.Z., P.D.K., B.S.G., J.R.M., D.C.M, M.A., G.D.S., K.W., K.O.S., P.A., B.F.H., R.S.B.; Wrote the first draft of the paper: D.R.M; Read and edited the paper: D.R.M, A.S., S.G., D.L., T.Z., P.D.K., B.S.G., J.R.M., D.C.M, M.A., G.D.S., K.W., K.O.S., P.A., B.F.H., R.S.B. Funding acquisition: D.R.M., G.D.S., B.F.H., R.S.B. All authors reviewed and approved the manuscript.

## Supplementary Figure Legends

**Figure S1.**
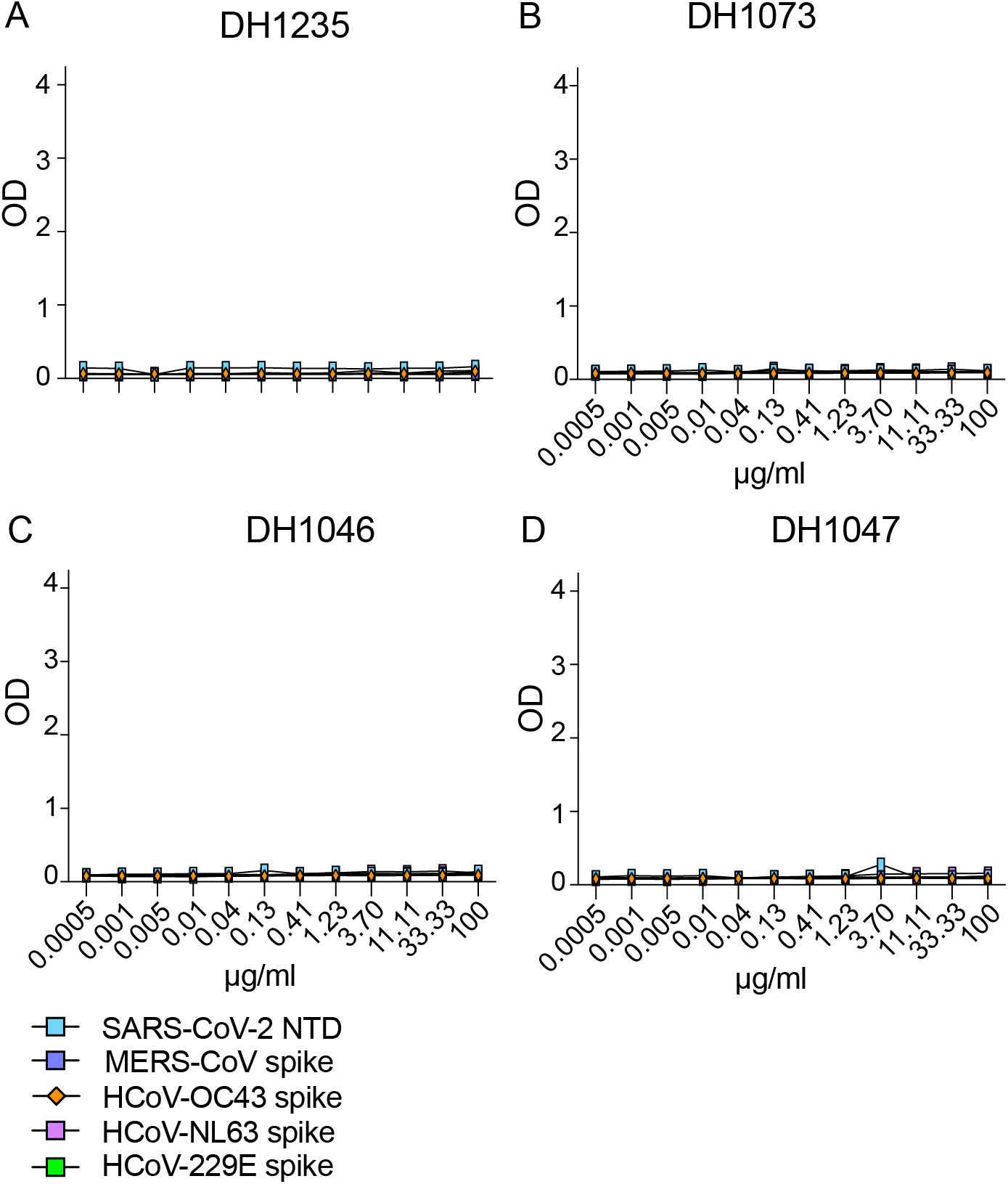
The binding activity of cross-reactive antibodies against MERS-CoV and human common-cold CoVs. The neutralization activity of four broadly neutralizing antibodies against SARS-CoV-2 NTD, MERS-CoV spike, HCoV-OC43 spike, HCoV-NL63, and HCoV-229E shown for (A) DH1235, (B) DH1073, (C) DH1046, and (D) DH1047.

**Figure S2.**
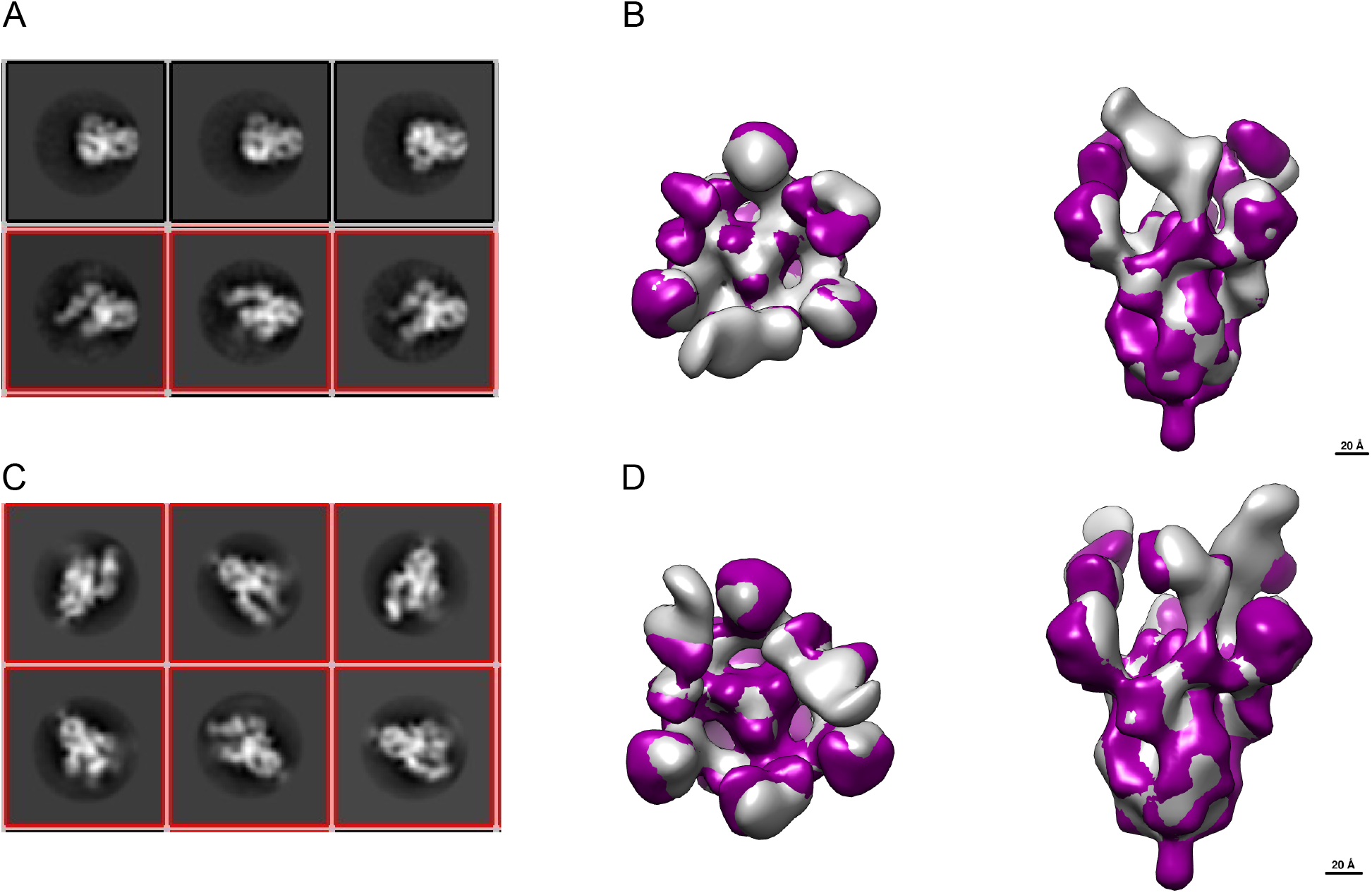
NSEM of DH1047 bound to bat RsSHC014 and SARS-CoV spike ectodomains. (A) Representative 2D class averages of bat RsSHC014 2P spike ectodomain bound to DH1047 Fab.
(B) Overlay of 3D reconstruction of DH1047 bound to bat RsSHC014 2P (grey) and SARS-CoV-2 HexaPro (purple) S ectodomains.
(C) Representative 2D class averages of SARS-CoV 2P spike ectodomain bound to DH1047 Fab (D) Overlay of 3D reconstruction of DH1047 bound to bat SARS-CoV 2P (grey) and SARS-CoV-2 HexaPro (purple) S ectodomains. The red boxes in panels A and C indicate the classes that show DH1047 Fab bound to spike.

**Figure S3.**
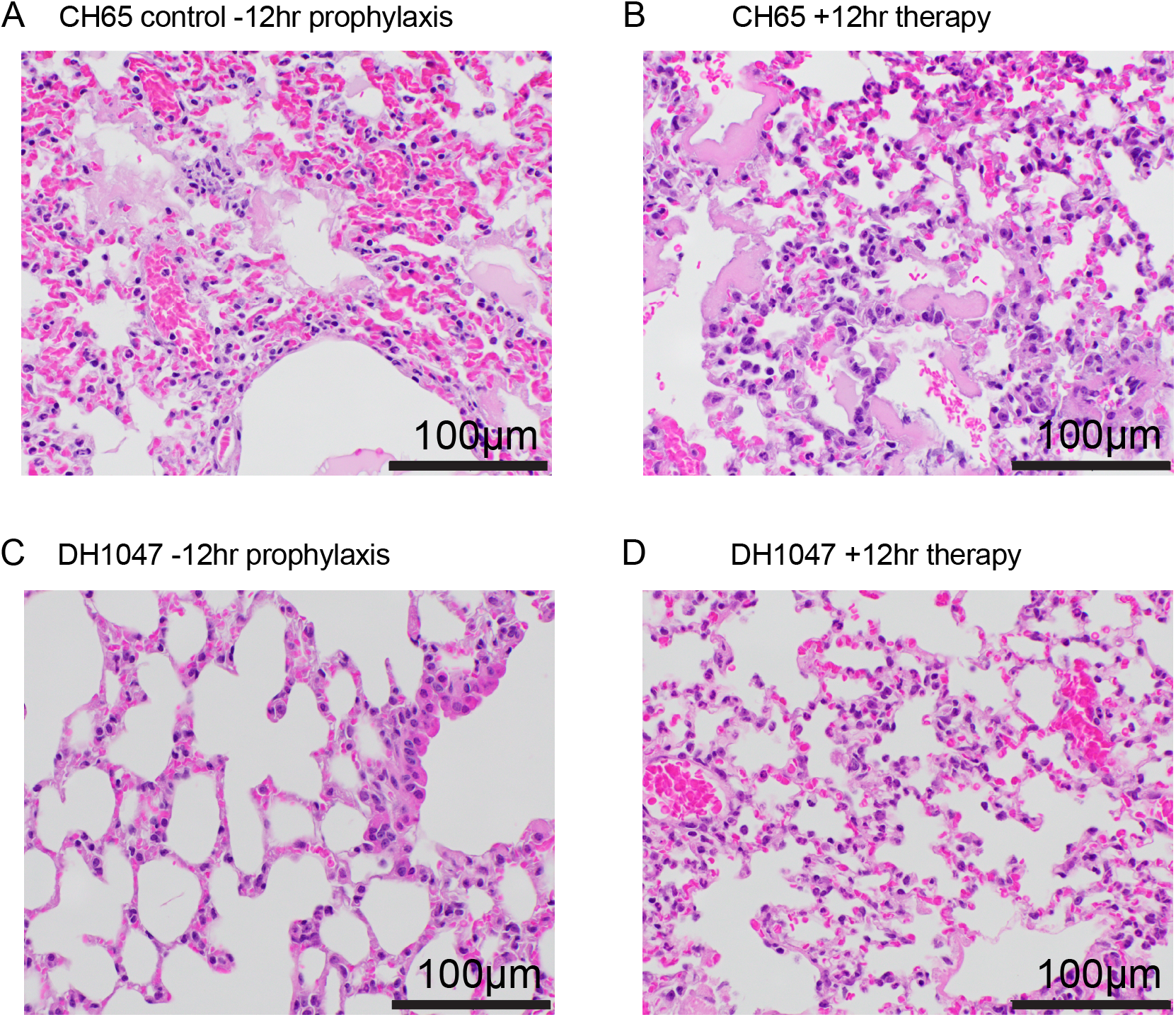
Lung H+E staining of SARS-CoV infected mice. Pathologic features of acute lung injury were scored using two separate tools: the American Thoracic Society Lung Injury Scoring (ATS ALI) system. Using this ATS ALI system, we created an aggregate score for the following features: neutrophils in the alveolar and interstitial space, hyaline membranes, proteinaceous debris filling the air spaces, and alveolar septal thickening. Three randomly chosen high power (×60) fields of diseased lung were assessed per mouse. Representative images are shown from vehicle and RDV-treated mice. All images were taken at the same magnification. The black bar indicates 100 μm scale. (A) CH65 control prophylaxis. (B) CH65 therapy. (C) DH1047 prophylaxis. (D) DH1047 therapy.

**Figure S4.**
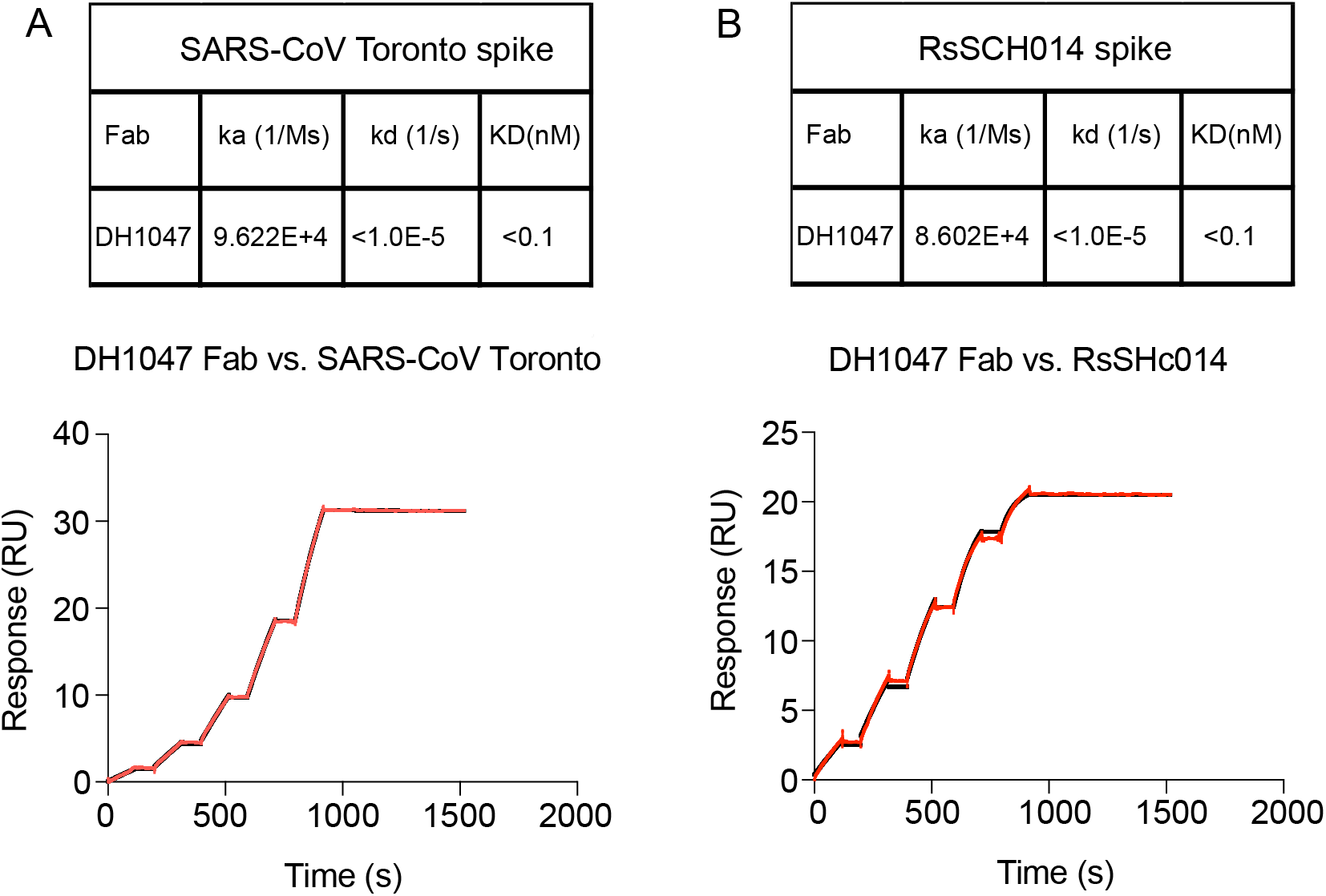
The affinity data of DH1047 against SARS-CoV and RsSHC014 spikes. Surface plasmon resonance (SPR) binding experiments of DH1047 against (A) SARS-CoV-2 Toronto and (B) RsSHC014. Binding affinity measurements are shown in the tables and response units (RU) as a function of time in seconds (s) is shown for both SARS-CoV and RsSHC014. SPR experiments were repeated twice.

**Figure S5.**
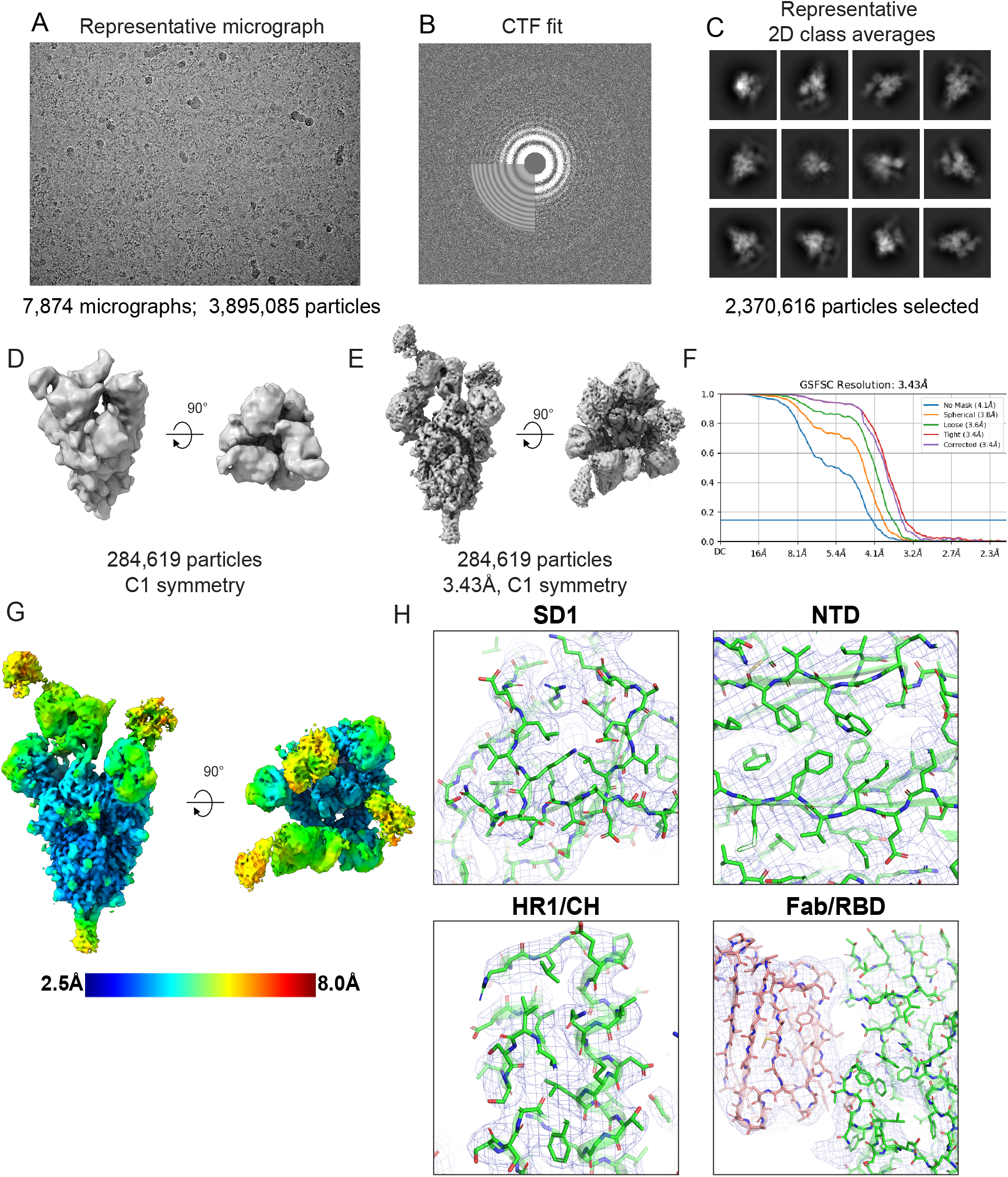
Cryo-EM data processing for the SARS-CoV spike ectodomain bound to DH1047, Related to Figure 2. (A) Representative cryo-EM micrograph.
(B) Cryo-EM CTF fit.
(C) Representative 2D class averages from Cryo-EM dataset.
(D) *Ab initio* reconstruction.
(E) Refined map.
(F) Fourier shell correlation curve.
(G) Refined cryo-EM map colored by local resolution.
(H) Zoom-in images showing the SD1, NTD, HR1/CH and RBD/Fab contact regions in the structure. The cryo-EM map is shown as a blue mesh and the fitted model is in cartoon representation, with residues shown as stick.

**Figure S6.**
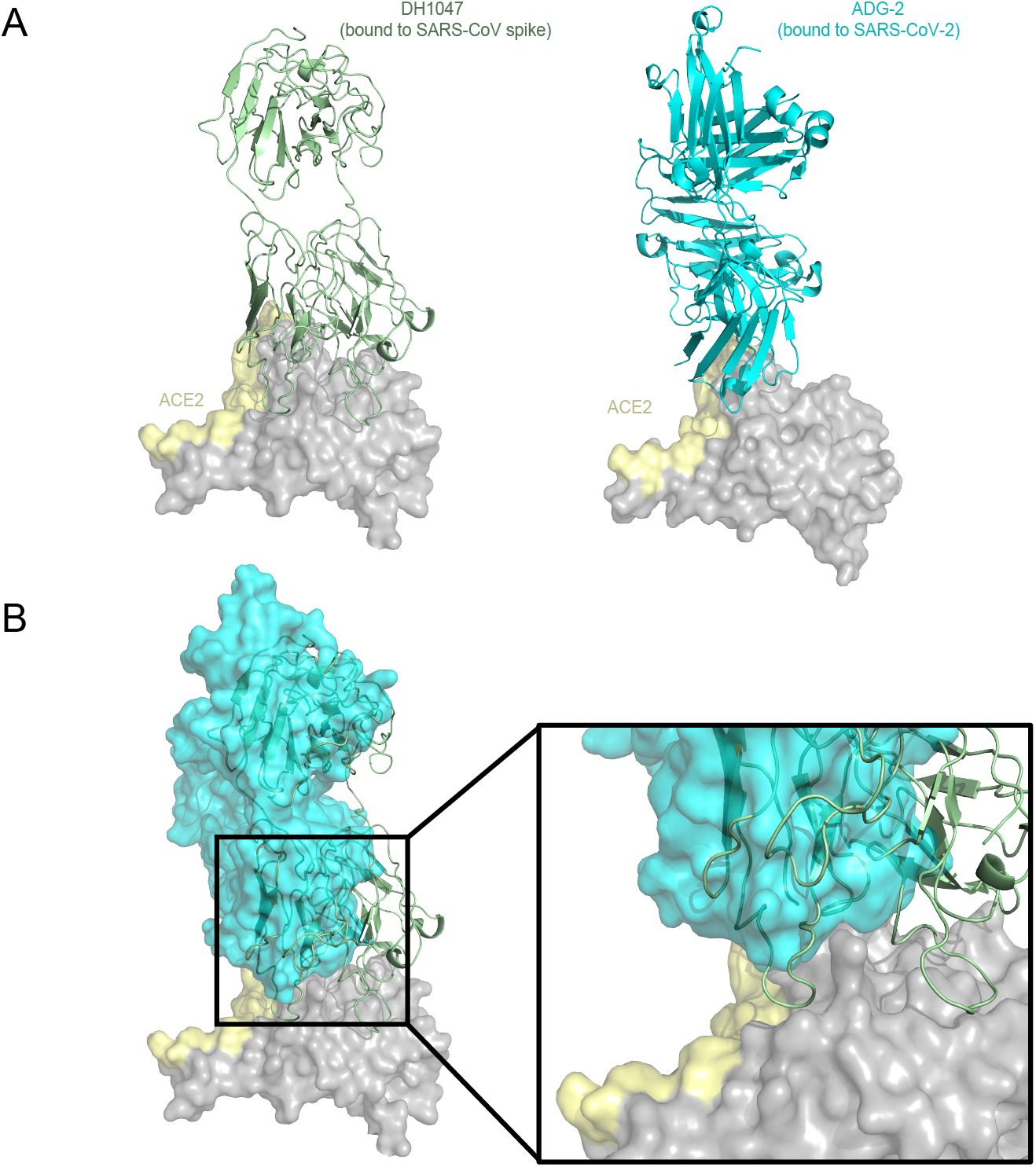
DH1047 and ADG-2 binds the RBD of SARS-Cov and SARS-CoV-2 spike ectodomains using a similar footprint. (A) Cartoon representation of DH1047 (colored in pale green) bound to the RBD (grey surface, ACE2 binding site in yellow) of SARS-CoV S ectodomain and ADG-2 (cyan) bound to SARS-CoV-2 S ectodomain. The homologous Fab ADI-19425 (PDB 6APC) was docked in the ADG-2 cryo-EM map (EMD-23160) to generate the model.
(B) DH1047 and ADG-2 bind partially overlapping binding sites on the RBD.

**Supplemental Table 1:**
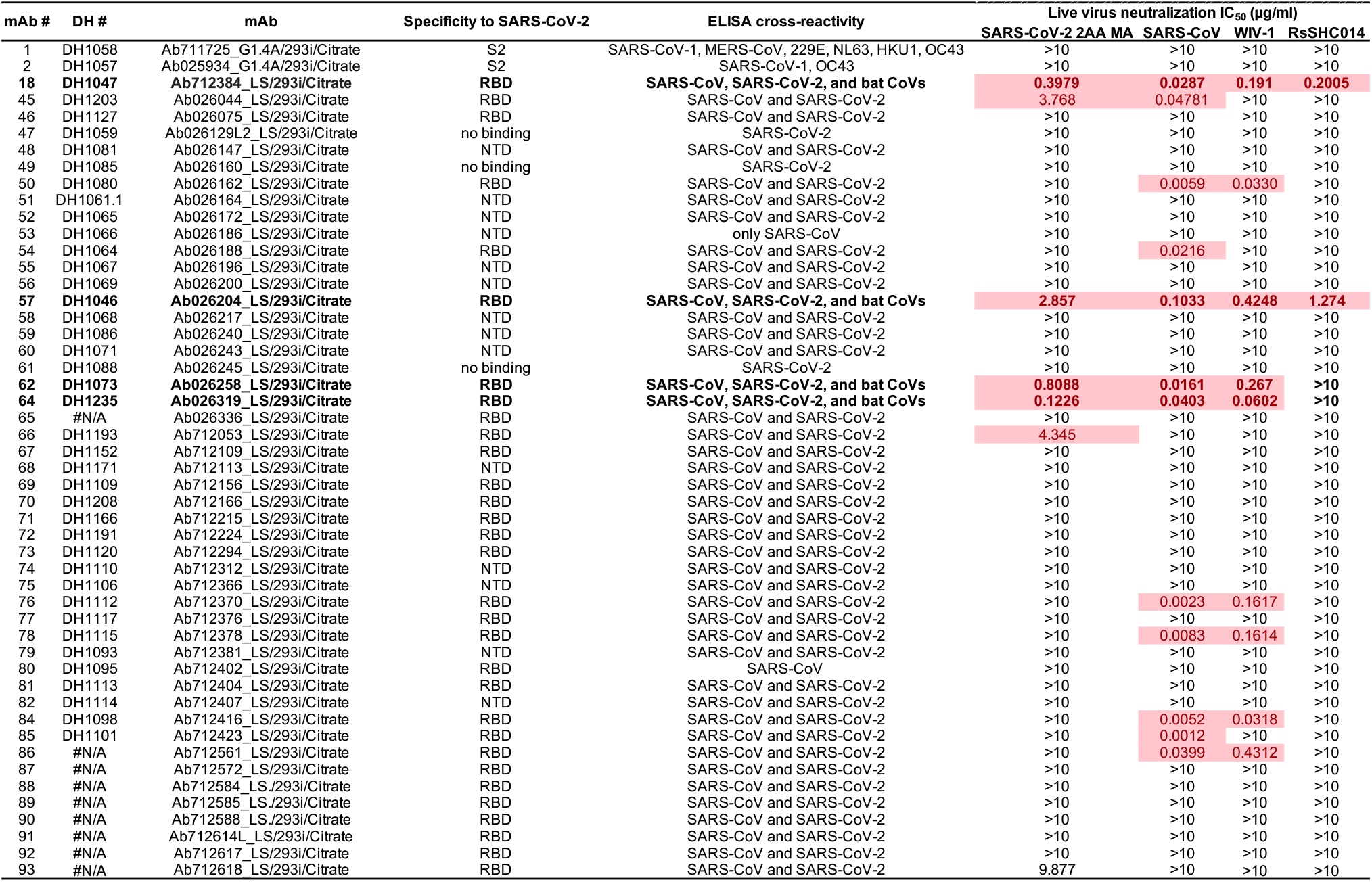
monoclonal antibody screen against SARS-CoV-2 2AA MA, SARS-CoV, WIV-1, and RsSHC014.

**Supplemental Table 2:** Immunogenetic characteristics of broadly cross-reactive mAbs.

| DH# | Antibody ID | Binding Specificity | Cross Reactivity | Antibody Gene Analysis |  |  |  |  |  |  |  |  |  |
| --- | --- | --- | --- | --- | --- | --- | --- | --- | --- | --- | --- | --- | --- |
|  |  |  |  | Donor ID | Time Point | HCDR3 Length | Heavy chain mutation | VH_Gene | JH_Gene | LCDR3 Length | Light chain mutation | VL_Gene | JL_Gene |
| DH1235 | Ab026319_LS | RBD | SARS-CoV-1 | SARS-CoV-2 convalescent | Day 36 | 21 | 1.68 | IGHV3-48 | IGHJ4 | 9 | 1.75 | IGLV4-60 | IGLJ2 |
| DH1073 | Ab026258_LS | RBD | SARS-CoV-1 | SARS-CoV convalescent | Year 17 | 15 | 9.06 | IGHV1-46 | IGHJ6 | 11 | 2.92 | IGKV3-11 | IGKJ1 |
| DH1046 | Ab026204_LS | RBD | SARS-CoV, PCoV GXP4L, Bat CoV RsSHC014, Bat CoV RaTG13 | SARS-CoV convalescent | Year 17 | 24 | 4.70 | IGHV3-23 | IGHJ6 | 9 | 3.65 | IGKV1-5 | IGKJ2 |
| DH1047 | Ab712384_LS | RBD | SARS-CoV, PCoV GXP4L, Bat CoV RsSHC014, Bat CoV RaTG13 | SARS-CoV convalescent | Year 17 | 24 | 8.05 | IGHV1-46 | IGHJ4 | 9 | 2.05 | IGKV4-1 | IGKJ1 |

**Supplementary Table 3.**
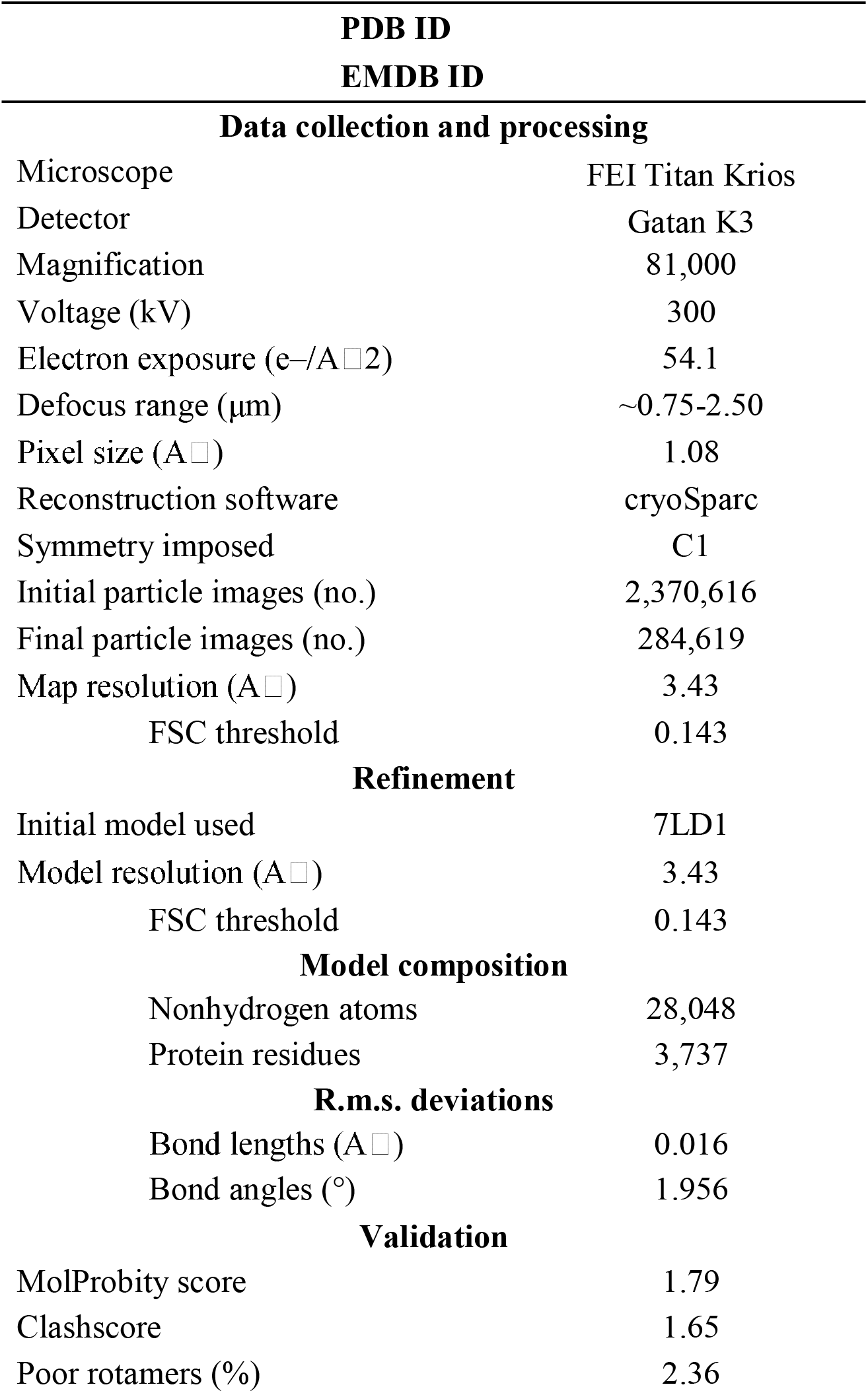
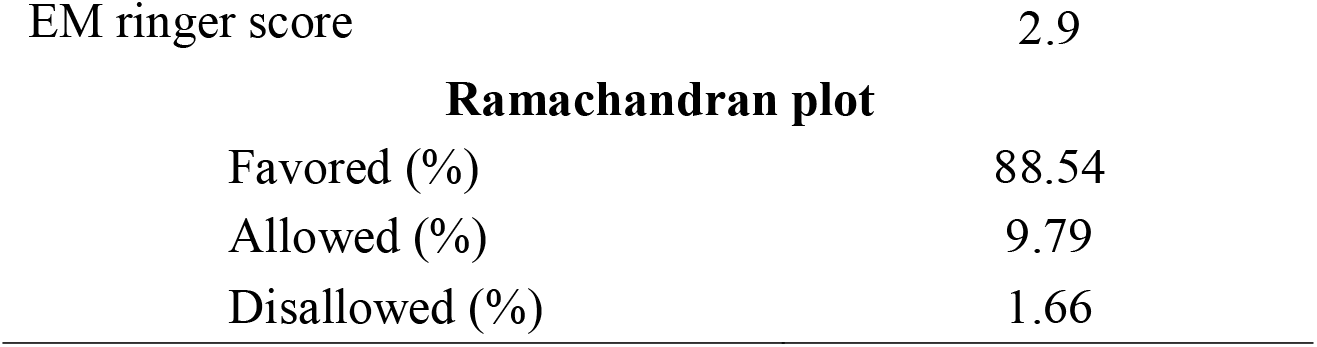
Cryo-EM data collection and refinements statistics.

